# Retinal waves shape starburst amacrine cell dendrite development through a direction-selective dendritic computation

**DOI:** 10.64898/2026.02.02.701812

**Authors:** Miah N. Pitcher, Aanica S.B. Gonzales, Raul Habib, Marla Feller

## Abstract

During development, dendrites undergo structural plasticity in response to neural activity; however, whether spatiotemporal activity patterns can instruct dendritic growth remains unclear. Prior to vision, the developing mouse retina exhibits spontaneous retinal waves with a nasal propagation bias that mimics forward optic flow. Here, we reveal that starburst amacrine cells use direction-selective dendritic computations to transform this propagation bias into asymmetric dendrite growth, linking activity patterns to structural development.

## INTRODUCTION

In the adult nervous system, dendritic computations distinguish spatiotemporal patterns of inputs. Dendrites filter, enhance, or suppress incoming synaptic signals through an interplay of passive, active, and morphological properties^1^. In addition, dendrite properties can selectively enhance certain spatiotemporal sequences of input^2–4^. During development, maturation of dendrite structure and synaptic function— including synapse distributions, postsynaptic strength, and dendrite morphology— combine to establish dendritic computations^5^. However, whether dendritic computations can in turn influence the development of dendrite morphology remains an open question.

Here, we use the mouse retina as a model system to test the hypothesis that a local dendritic computation decodes propagation direction of spontaneous activity during development to shape dendrite morphology. In the retina, starburst amacrine cells (SACs) provide a canonical example of a direction-selective dendritic computation^3^. SACs are axonless inhibitory interneurons with dendrites that extend radially around the soma^6–8^. Each dendrite receives excitatory input on the proximal dendrite and releases neurotransmitter from varicosities at the distal tip^9–11^. When a visual stimulus within a SAC’s receptive field moves in the centrifugal direction—from the soma toward the distal tip of an individual dendrite—the varicosities show a larger calcium transient compared to motion in the opposite direction^12^. Thus, individual dendrites act as compartmentalized local motion detectors that summate excitatory inputs to elicit a strong centrifugal motion response^10,11,13–17^.

SACs reach morphological maturity during the first two postnatal weeks, before eye-opening^18^. During this time, the primary source of activity in the retina is spontaneous retinal waves. Early in the second postnatal week, retinal waves exhibit a bias to propagate in the nasal direction^19–23^, resembling the activity pattern of forward optic flow on the surface of the retina^19^. Propagation bias of waves has been proposed to instruct both direction-selective^19^ and retinotopic^24^ map formation, yet how cells in the retina read out propagation bias remains unknown. Here, we used two-photon calcium imaging to show that developing SAC dendrites exhibit direction-selective responses to retinal waves. Using mouse models with altered retinal waves, we show that this local direction-selective response drives asymmetric dendritic growth during propagation bias, indicating that wave direction plays an instructive role in morphological development of SAC dendrites.

## RESULTS

### SAC dendrites compute light and wave direction prior to eye opening

We began by determining the age at which SACs first exhibit direction-selective dendritic computations using two-photon calcium imaging combined with light stimulation. SACs first showed consistent dendrite calcium responses to moving bars at P12, prior to eye-opening. Moreover, individual SAC dendrites in mice P12-13 show a higher response to bars moving in centrifugal (CF) directions than centripetal (CP) directions, with tuning properties comparable to adult (Fig 1A-C, Fig S1). Hence, the direction-selective computation in SAC dendrites is present at the onset of light responsiveness in the retina.

**Figure 1.**
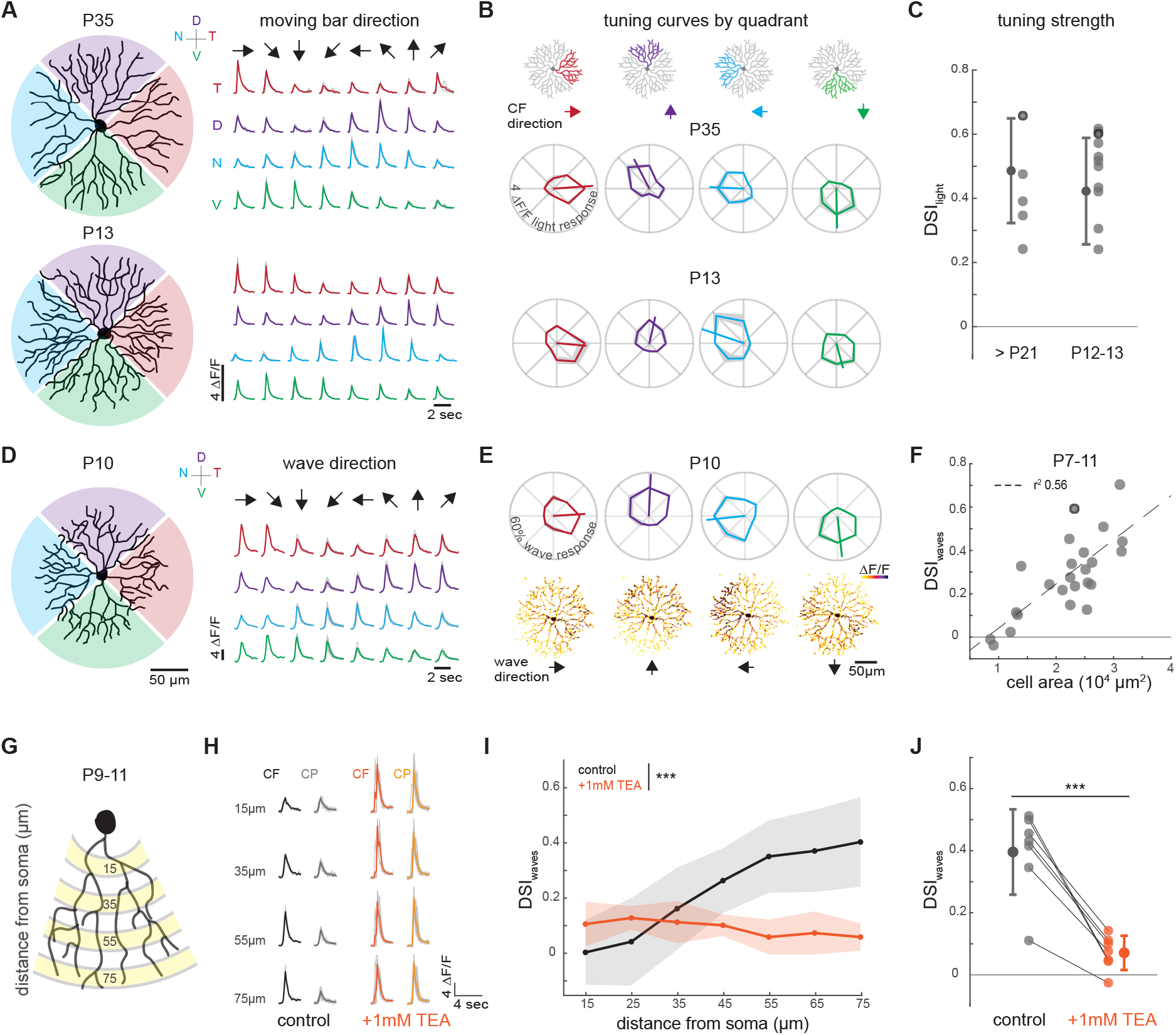
SAC dendrites compute light and wave direction prior to eye opening. (A) Reconstruction of SAC from adult (top) and P13 (bottom). Traces are from example cells on the left and show average ΔF/F_0_ response of each quadrant to three repetitions of moving bars in eight directions. Grey lines show individual trials; colors indicate quadrant orientation (red = temporal, blue = nasal, purple = dorsal, green = ventral). Black arrows indicate direction of moving bars. (B) Tuning curves for each quadrant of example cells in (A). Lines show vector sum of average response, shaded regions show standard deviation across three trials. Centrifugal (CF) direction for each quadrant indicated by the colored arrow, centripetal (CP) direction is 180° opposite. (C) Tuning strength (DSI_light_ = normalized difference between quadrant response to bars moving in centrifugal directions vs centripetal directions, see Methods) for adult cells (n = 5) and cells P12-13 (n = 10). Each cell tuning value is the average of all four of its quadrants. Example cells outlined in black. No difference in DSI_light_ found at P12-13 compared to adult (two-sample t test, p = 0.3951). See also Fig S1. (D) Reconstruction of SAC at P10. Traces show average ΔF/F_0_ response of each quadrant (colored lines) to retinal waves propagating in each direction. Grey lines show individual wave responses; black arrows indicate wave direction. (E) Top: tuning curves for each quadrant of example cell in D. Polar axis represents normalized wave response (see Methods). Lines show vector sum of average response; shaded regions show standard deviation. Bottom: normalized ΔF/F_0_ response of the same cell to waves in four different directions indicated by black arrows. Darker colors represent higher responses. (F) Scatter plot showing cell tuning strength (DSI_wave_ represents normalized difference between quadrant response to waves propagating in centrifugal directions vs centripetal directions) as a function of cell convex hull area, with a linear regression fit. Cell area significantly predicted tuning strength (n = 24 cells, linear regression, R^2^ = 0.56, p = 2.58e-05). Example cell outlined in black. (G) Schematic showing segmentation of quadrant into ROIs of 10 µm width rings extending outward from the soma to test for evidence of summation. Numbers indicate distance between the center of the ring and the soma. Example responses from ROIs highlighted in yellow are shown in h. (H) Left: Calcium transients from one example quadrant showing the response to waves going in the centrifugal direction (black) and centripetal direction (grey). Right: responses of the same ROIs after application of 1mM TEA to waves going in the centrifugal (dark orange) and centripetal (light orange) direction. (I) Summary data across P9-11 cells (n = 7) showing tuning strength (DSI_wave_) as a function of distance from the soma in control (grey) and after application of TEA (orange). Linear mixed-effects modeling showed a strong drug-by-distance interaction (p = 9.8e-16), such that tuning strength increased with distance from the soma in control conditions but was independent of distance following TEA application. (J) Average tuning strength by cell shows a decrease after application of TEA (paired t-test, p=2.558e-4). All data are presented as mean values ± standard deviation.

Next, we tested whether SAC dendrites show direction-selective responses to retinal waves earlier in development. We used dual-colored two-photon imaging to simultaneously measure wave direction and SAC dendrite calcium responses in mice aged P7-P11. By P10, individual SAC dendrites showed a higher response to waves propagating in CF than the CP directions (Fig 1D-E). We measured the strength of tuning to wave direction of each quadrant using a direction selective index (DSI_wave_), defined as the normalized difference in response to CF- and CP-propagating waves (see Methods). Across cells aged P7-P11, tuning strength varied substantially and was correlated with cell size, with larger cells exhibiting stronger tuning (Fig 1F). In mature SACs, direction-selectivity increases with distance from the soma due to summation of excitatory inputs along electrically compartmentalized dendrites^10,13–15^ (Fig S1). We therefore hypothesized that, during development, SAC dendrites decode wave direction using a similar computation, such that increased dendritic length enhances summation and tuning strength.

To test this hypothesis, we first examined whether tuning strength increased along the dendrite, consistent with spatiotemporal summation of excitatory inputs in the preferred direction. In P9-11 SACs, tuning strength increased as a function of distance from the soma, due to selective enhancement of the centrifugal wave response (Fig 1G, H). Next, we tested whether disrupting dendritic compartmentalization compromised the directional tuning of dendrites. Mature SAC dendrites form independent computational units due to the presence of a large conductance, potassium channel Kv3.1, that shunts depolarizations near the soma^13,15^. After bath application of tetraethylammonium (TEA), a blocker of multiple K+ channels including Kv3.1, SAC dendrites responded equally to all waves regardless of propagation direction, leading to a decrease in tuning strength across the dendrite (Fig 1H-J). This change in tuning was not explained by changes in the directional propagation of waves after drug application, nor by saturation of the calcium sensor (Fig S2). Together, these results demonstrate that SAC dendrites decode wave direction through early maturation of summation and electrical compartmentalization, enabling a localized dendritic direction-selective computation.

### Retinal wave propagation bias instructs SAC dendrite growth

From P8-P11, retinal waves exhibit a bias to propagate toward nasal retina^19–23^. Our results demonstrating that SAC dendrites show direction-selective responses to waves predict that dendrites oriented in the nasal direction will be preferentially activated during propagation bias. SAC dendrites are also growing rapidly during waves, showing an approximately fivefold expansion in dendrite territory in the first two postnatal weeks^18^. We hypothesized that 1) activity from waves promotes SAC dendrite growth and 2) during propagation bias, preferential activation of the nasal-oriented dendrites accelerates nasal dendrite growth.

First, we determined that SAC dendrites exhibit activity-dependent dendrite growth. During the first postnatal week, SACs are depolarized by retinal waves mediated by cholinergic signaling (Fig 2A). Cholinergic waves are severely disrupted in a mouse model in which the β2 subunit of the nicotinic acetylcholine receptor is genetically knocked out (β2-nAChR-KO)^25,26^. We found a significant decrease in spontaneous calcium transients in SACs at P8 in β2-nAChR-KO mice (Fig 2C-D). We also observed a significant decrease in total dendrite length and complexity in P8 β2-nAChR-KO SACs compared to littermate controls (Fig 2E-G). Previous reports have shown no difference in dendritic arbor size in adult β2-nAChR-KO SACs^23^, indicating that chronic absence of the β2-nAChR receptor itself does not prevent growth and suggests instead an acute effect of activity loss on growth (as previously observed in many developing dendrites^27–30^). Together, these results show that wave-induced activity in SACs during the first postnatal week promotes dendrite growth.

**Figure 2.**
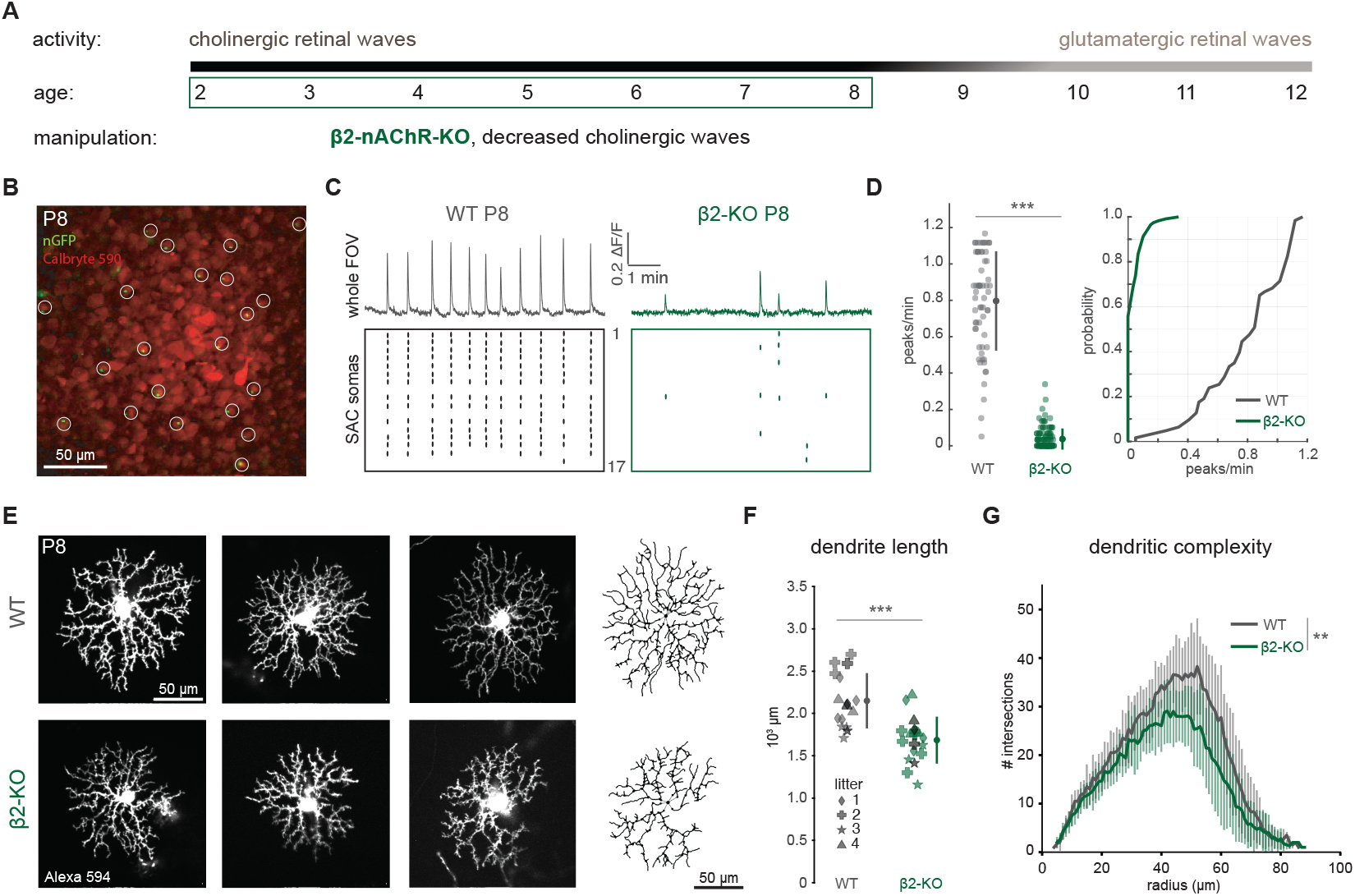
Retinal waves promote SAC dendrite growth during first postnatal week. (A) Timeline of spontaneous activity during development. Retinal waves are mediated by cholinergic signaling in the first postnatal week and are significantly perturbed in β2-nAChR-KO mice. (B) Example image of retina expressing nGFP in the nucleus of SACs (Chat.Cre/nGFP) bolus loaded with Calbryte^TM^ 590 AM. White circles show SAC soma ROIs identified by nGFP expression. (C) ΔF/F_0_ calcium traces of waves (top) and raster plot of SAC activity (bottom) in P8 wild-type and β2-nAChR-KO mice. (D) Calcium transient frequency in wild-type SACs at P8 (n = 63 cells across 4 FOVs in 3 retinas) and β2-nAChR-KO SACs (n = 115 cells across 6 FOVs in 4 retinas). β2-nAChR-KO SACs show a significant reduction in activity (Mann–Whitney U test p = 5.928e-29). Cumulative distribution function shown on the right. (E) Example images of SAC dendrites filled with Alexa 594 in wild-type (top row) and β2-nAChR-KO (bottom row). Columns indicate littermate pairs. Right panel shows skeleton tracings from SNT. (F) Total dendritic length of wild-type (n = 12 cells from 4 mice) and β2-nAChR-KO (n = 15 cells from 4 mice) SACs. Different symbols denote cells from littermate pairs; darker symbols indicate litter averages. β2-nAChR-KO SACs show a significant reduction in path length (two-sample t-test, p = 0.0005). (G) Sholl analysis comparing dendritic complexity in wild-type and β2-nAChR-KO SACs. β2-nAChR-KO SAC dendrites show reduced overall complexity with uniform downward shift (linear mixed effects model found significant main effect of genotype (β = –3.50, p = 0.0024) and no genotype × radius interaction (p = 0.40)). All data are presented as mean values ± standard deviation

Next, we tested whether propagation bias affects the activation and maturation of the nasal quadrant (Fig 3A). In wild-type mice, we observed a strong propagation bias in waves from P8-P11 (Fig 3B), which led to increased activity in the nasal quadrant compared to the temporal quadrant during 30 mins of waves (Fig 3C). Further, in SACs from mice P9-P11—after as little as one day of propagation bias—nasal quadrants exhibited higher dendritic growth than temporal quadrants (Fig 3F), both in terms of total dendrite length (Fig 3G), and distal dendrite complexity (Fig 3H).

**Figure 3.**
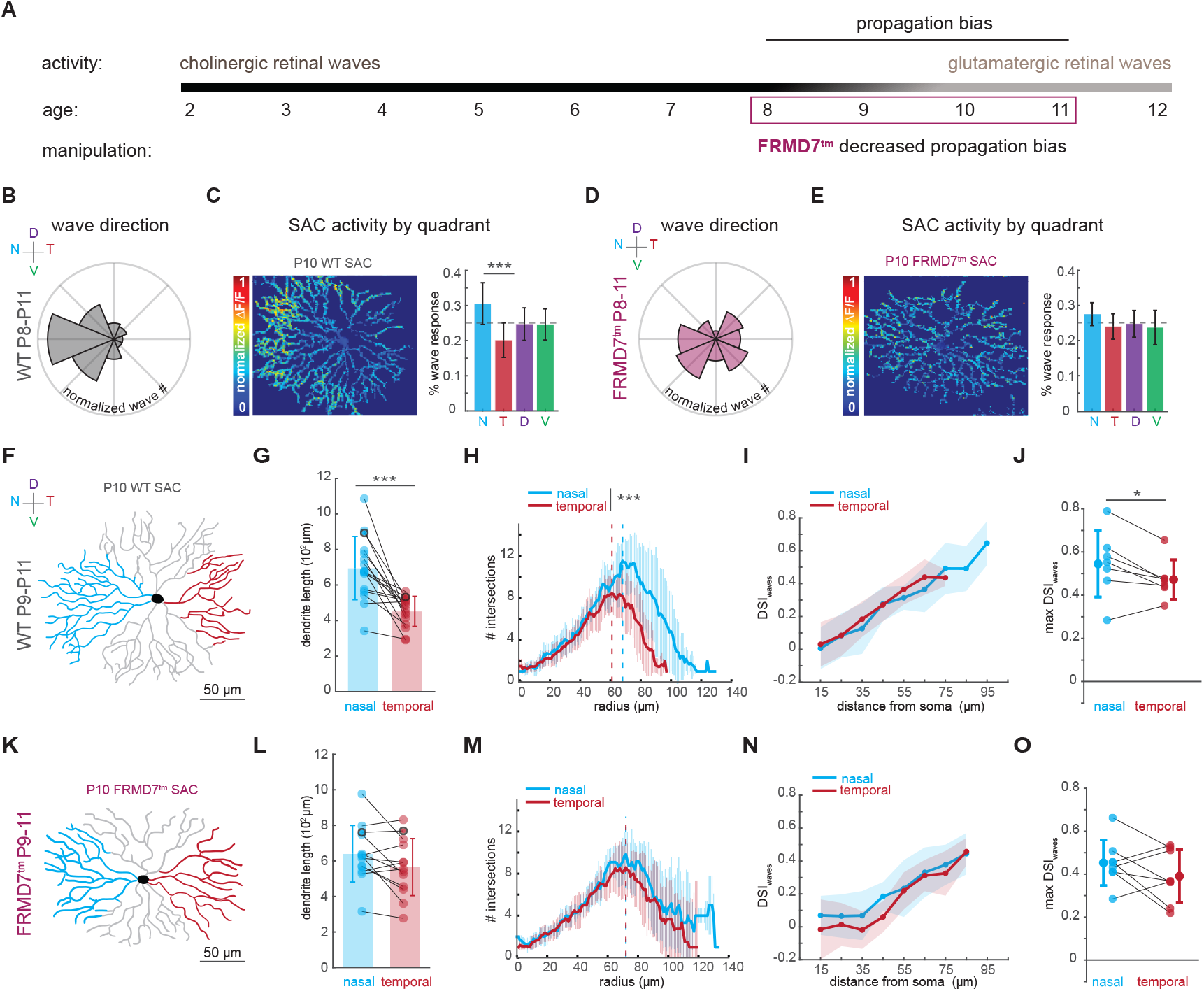
Retinal wave propagation bias instructs SAC dendrite growth. (A) Timeline of spontaneous activity during development. Retinal waves show propagation bias from P8-P11 which is decreased in FRMD7^tm^ mice. (B) Distribution of retinal wave propagation direction in wild-type mice aged P8-P11 (n = 158 waves from 6 FOVs in 4 retinas). (C) SAC dendrite activity in response to retinal waves during propagation bias. Left: cumulative activity (normalized sum of ΔF/F_0_) in a wild-type SAC at P10 in response to 30 minutes of retinal waves. Right: summary data showing the normalized sum of wave response in each quadrant across 30 minutes of retinal waves (n = 12 cells). Nasal quadrant wave response is increased compared to temporal quadrant (p = 8.9e-4, mixed-design ANOVA on CLR transformed data followed by Tukey’s HSD post-hoc, see Methods). (D) Distribution of retinal wave propagation direction in FRMD7^tm^ mice aged P9-11 (n = 348 waves from 10 FOVs in 4 retinas) (E) Same as (C) but for FRMD7^tm^ SACs (n = 10 cells). No significant differences between quadrants were detected in FRMD7 cells (all p > 0.40, mixed-design ANOVA on CLR transformed data followed by Tukey’s HSD post-hoc, see Methods). (F) Example P10 SAC with dendrites from nasal and temporal quadrants pseudo colored in blue and red, respectively. (G) Summary data showing path length of nasal and temporal quadrants in wild-type (n = 15 cells) SACs. Example cell in f outlined in black. SACs show higher path length in nasal compared to temporal dendrites (p = 1.5e-6, mixed-design ANOVA followed by Tukey’s HSD post-hoc). (H) Sholl analysis across nasal and temporal quadrants in wild-type SACs. Dashed lines indicate Sholl peaks. Wild-type cells show increased density in nasal compared to temporal distal dendrites (linear mixed effects model found genotype × quadrant × radius interaction: β = 0.014 ± 0.006, p = 0.012, with bin-based mixed-effects analysis showing greater nasal–temporal asymmetry in dendrites > 60 µm from soma: β = 2.07 ± 0.71, p = 0.0045). (I) Tuning strength (DSI_wave_) as a function of distance from soma across nasal and temporal dendrites in wild-type SACs (n = 7 cells). (J) Peak tuning strength (DSI_wave_) increased in nasal compared to temporal dendrites for wild-type SACs (n= 7 cells, p = 0.0468, paired t-test) (K-M) Same as (F-H) for FMRD7^tm^ SACs (n = 13 cells). No difference found between nasal and temporal path length (p = 0.0999, mixed design ANOVA followed Tukey’s HSD post-hoc), and asymmetry in density was significantly reduced relative to wild-type (linear mixed effects model, significant genotype × quadrant interaction (β = –1.00 ± 0.33, p = 0.0027)) (N-O) Same as (I-J) for FRMD7^tm^ SACs (n = 8 cells). No difference found in peak tuning between nasal and temporal dendrites (p = 0.21, paired t-test). Data are presented as mean values ± standard deviation.

We next aimed to alter wave propagation direction and test whether the dendrite asymmetry was disrupted. Although the mechanisms underlying propagation bias remain incompletely understood^19,23,31^, a mouse model with a mutation in the gene *FRMD7* (FRMD7^tm^)^32^ has been previously shown to exhibit reduced propagation bias^19^, a result that we replicated (Fig 3D). We found that dendrites in FRMD7^tm^ SACs compute wave direction (Fig S4) and show an equal activation of nasal and temporal quadrants during 30 mins of waves (Fig 3E). We reconstructed SACs from FRMD7^tm^ mice P9-P11 and found no difference in dendritic length in nasal and temporal dendrites (Fig 3K, L), with significantly less asymmetry in dendrite density as compared to wild-type (Fig 3M). These experiments do not exclude FRMD7-intrinsic effects on SAC morphology that are independent of wave propagation. However, the spatial and temporal profiles of the effect are inconsistent with a cell-intrinsic mechanism: FRMD7 is symmetrically localized in SACs^32^, and the morphological asymmetry we find is not yet present at P8 (Fig S3) when FRMD7 expression is already high^32^. Instead, this effect only emerges at P9-11 after propagation bias has begun.

Lastly, we investigated whether asymmetry in dendrite morphology contributes to asymmetry in dendrite tuning properties across the horizontal axis. In wild-type SACs, both nasal and temporal dendrites show increased tuning strength as a function of distance from the soma (Fig 3I), but the nasal dendrites showed a higher peak tuning strength due to their increased outgrowth (Fig 3J). In FRMD7^tm^ SACs, there was no difference in peak tuning strength between nasal and temporal dendrites (Fig 3N, O). While the wild-type SAC asymmetry in tuning strength is modest, it could reinforce asymmetric activity along the horizontal axis and promote further differential growth through positive feedback. Together, these results suggest that SAC dendrite’s direction-selective responses to retinal waves can convert biases in wave propagation into asymmetric dendritic maturation.

## DISCUSSION

In this study, we describe an unexpected developmental role for starburst amacrine cell (SAC) dendrites as active computational units. We show that SAC dendrites exhibit direction-selective calcium responses to both visual motion at the onset of light responsiveness and, earlier, to the propagation direction of spontaneous retinal waves. This direction-selective response depends on spatiotemporal summation and electrical compartmentalization—core features of mature SAC function—indicating that a canonical direction-selective dendritic computation is established early in development. Crucially, this computation enables SACs to read out biases in wave propagation and convert them into asymmetric dendritic growth, preferentially promoting outgrowth of dendrites aligned with the direction of wave propagation bias. Together, our findings demonstrate how early dendritic computations transform patterned spontaneous activity into structural asymmetries that shape developing neural circuits.

The instructive role of spatiotemporal activity patterns in circuit development has been well established in the visual system. Retinal waves drive correlated firing among neighboring retinal ganglion cells while decorrelating activity among distant cells, embedding information about spatial relationships into neural activity^33^. Manipulations that alter the spatiotemporal structure of retinal waves reveal that distinct activity statistics instruct different features of visual map formation in downstream visual areas: local intraretinal correlation is important for retinotopic map formation, while anticorrelation between eyes is important for eye-specific segregation^26,34,35^. In each of these cases, Hebbian plasticity rules explain how activity shapes connectivity: correlated activity strengthens synapses, while uncorrelated activity weakens them, leading to competitive refinement^35^.

An instructive role for wave propagation bias has also been proposed, motivated by its similarity to the future optic flow experienced during self-motion. In *Xenopus* tadpoles, which lack retinal waves and instead experience optic flow induced by swimming during development, experimentally reversing the direction of visual motion disrupts retinotopic map formation in the optic tectum^24^. In mice, altering wave propagation bias reduces directional tuning in superior colliculus neurons^19^. Our results identify the earliest decoding of wave propagation bias as occurring via a dendritic computation in SACs, revealing an instructive mechanism that is distinct from classical Hebbian plasticity and operates through subcellular decoding of structured population activity.

Asymmetry in SAC dendrite maturation has important implications for the development of direction-selective circuits. In the mature retina, direction-selective ganglion cells (DSGCs) receive asymmetric inhibition from SACs that shapes their tuning. Each quadrant of a SAC preferentially forms inhibitory synapses with one DSGC subtype, tuning that DSGC to motion opposite the SAC quadrant orientation^36^. This precise synaptic wiring emerges during the second postnatal week^37–39^ and is disrupted in the mouse models used in this study with altered retinal waves^23,32^. Interestingly, these wave manipulations have axis-specific effects on DSGC tuning, showing altered synapse specificity with SACs along the horizontal, but not vertical axis^23,32,40^.

While the mechanisms that instruct synapse specificity in this circuit remain a mystery, our data demonstrate that SAC dendrites can use a dendritic computation in combination with retinal wave propagation bias to intrinsically differentiate between nasal and temporal quadrants during the second postnatal week, when these quadrants are synapsing onto distinct horizontal DSGC subtypes. This motivates future work exploring whether the differential activity or outgrowth across the horizontal axis of SAC dendrites instructs synaptogenesis with the correct postsynaptic partners.

Beyond these circuit-level implications, our findings expand the current framework of activity-dependent dendrite development. Compartmentalized calcium signaling in developing dendrites has been shown to locally influence dendritic structure in many regions of the developing central nervous system, with the prevailing model proposing that dendritic branches receiving afferent activity are selectively stabilized while inactive or uncorrelated branches are pruned^27–30,41–45^. Although the intracellular signaling pathways linking activity to dendritic growth are not fully understood, local elevations in intracellular calcium are thought to play a critical role^41,42^. Our results illustrate that compartment-specific calcium signals that regulate dendritic growth can arise not only from differences in input localization, but also from a dendritic computation that enhances specific spatiotemporal patterns of input. More broadly, this mechanism suggests that early maturation of dendritic computations may provide a strategy by which developing neurons translate structured network activity into spatially precise morphological refinement.

## RESOURCE AVAILABILITY

### Lead contact

Requests for further information and resources should be directed to and will be fulfilled by the lead contact, Marla Feller.

### Materials availability

This study did not generate new unique reagents.

### Data and code availability

Source data and analysis scripts supporting the findings of this study are available from the lead contact upon request.

## ACKNOWLEDGMENTS

All authors were supported by the National Institutes of Health (NIH) R01EY013528, R01EY019498, and P30EY003176. M.N.P was supported by NIH 1F31EY035571. We thank Ben Smith for his microscopy advice and support. We thank the members of the Feller, Puthussery, and Taylor labs, Yvette Fisher, Karthik Shekhar, Hillel Adesnik, and Dan Feldman for their comments on the manuscript.

## AUTHOR CONTRIBUTIONS

Conceptualization, M.N.P. and M.B.F.; methodology, M.N.P and M.B.F.; Investigation, M.N.P. and A.S.B.G., visualization M.N.P, A.S.B.G, and R.H., writing—original draft, M.N.P. and M.B.F; writing— review & editing, M.N.P. A.S.B.G, R.H and M.B.F.; funding acquisition, M.N.P and M.B.F.; supervision, M.B.F.

## METHODS

### ANIMALS

All animal procedures were approved by the UC Berkeley Institutional Animal Care and Use Committee and conformed to the NIH *Guide for the Care and Use of Laboratory Animals*, the Public Health Service Policy, and the SFN Policy on the Use of Animals in Neuroscience Research. Mice used in this study were aged from P7-P120 and were of both sexes. For most experiments, SACs were targeted using mice with Cre driven by the endogenous choline acetyltransferase promoter (Chat.Cre, Jackson Laboratories strain#:031661) crossed to mice with nuclear-localized GFP-lacZ function protein downstream of a loxP-flanked STOP sequence (nGFP, Jackson Laboratories strain #:008516). Chat-Cre/nGFP mice were then crossed with mice where the beta2 subunit of the nicotinic acetylcholine receptor is knocked out (β2-nAChR^−/−^ bred in lab). For SAC morphology development experiments, control animals were ChAT-Cre/nGFP β2-nAChR ^+/+^ or ChAT-Cre/nGFP β2-nAChR ^+/−^ littermates of ChAT-Cre/nGFP β2-nAChR ^−/−^ animals. FRMD7^tm^ mice refer to homozygous female or hemizygous male *Frmd7*^*tm1a(KOMP)Wtsi*^ mice (MMRRC at UC Davis, RRID:MMRRC_047759-UCD).

### METHOD DETAILS

#### Retina Preparation

Mice were deeply anesthetized with isoflurane inhalation and euthanized by decapitation. Eyes were enucleated and retinas were dissected under infrared light in oxygenated (95% O2/ 5% CO2) Ames’ media (Sigma) for light responses or ACSF (in mM, 119 NaCl, 2.5 KCl, 1.3 MgCl_2_, 1 K_2_HPO_4_, 26.2 NaHCO_3_, 11 D-glucose, and 2.5 CaCl_2_) for wave responses. Cuts along the choroid fissure were made prior to isolating the retina from the retinal pigmented epithelium to precisely orient retinas as described previously^23^. Isolated retinas were mounted whole over white filter paper (Whatman) with the ganglion cell layer facing up and dorsal retina oriented toward the top of the paper. Retinas were stored in room temperature oxygenated Ames or ACSF until use.

#### Two-photon calcium imaging

Two-photon fluorescence measurements were obtained with a modified movable objective microscope (MOM) (Sutter instruments, Novator, CA) and made using a Nikon 16X, 0.80 NA, N16×LWD-PF objective (Nikon, Tokyo, Japan). Two-photon excitation of fluorescent dyes was evoked with an ultrafast pulsed laser (Chameleon Ultra II; Coherent, Santa Clara, CA) tuned to 920 nm-1040 nm. The microscope system was controlled by ScanImage software (www.scanimage.org). This MOM was equipped with a through-the-objective light stimulation and two detection channels for fluorescence imaging.

#### SAC dendrite dye injections

SACs were filled with either 8 mM Calbryte™ 520, potassium salt (AAT Bioquest) for calcium imaging or 2 mM Alexa Fluor 594 (ThermoFisher) for morphological reconstructions via a sharp electrode as previously described^11,46^. Briefly, borosilicate glass (I.D. = 1.1, O.D. = 1.5 mm) electrodes were pulled to a resistance of 100-150 MΩ then bent (such that electrode tips were perpendicular to the retinal surface when inserted into the headstage). Bent electrodes were filled with dye and placed above the targeted SAC under infrared illumination. In cells with nGFP in SACs, two-photon illumination was used to guide the electrode tip through the inner limiting membrane into the targeted SAC and visualize cell filling. Otherwise, SACs were targeted using epifluorescence by soma shape and size and confirmed via dendritic morphology after the fill. Ionotophoresis of dye was achieved with negative current pulses (−10 to −20 nA, 500 ms) and electrodes were withdrawn as cell bodies began to fill. Filled SACs were left to recover for 30-60 minutes before imaging. All functional recordings were done at ∼32 °C.

#### SAC light responses

For light stimulation experiments, SACs in ventral retina were injected with calcium dye as described above and presented with visual stimuli. Stimuli patterns were generated in MATLAB using the psychophysics toolbox and projected onto the retina using a digital micromirror device containing an LED (UV: 375nm). Visual stimuli were delivered on the flyback of the fast axis scanning mirror during a unidirectional scan to interleave stimulus with imaging and decrease background signal from UV stimulation in the photomultiplier tubes. Calcium responses from the entire SAC dendritic field were imaged using a 920 nm excitation wavelength and images were acquired at 5.92 Hz (128 × 128 pixels, 1ms/line, bidirectional scanning). Visual stimuli consisted of three trials of moving bars (25 × 500 μm) moving at 500 μm/sec in eight pseudorandomized directions parallel to the shorter axis of the bar across the entire stimulus plane.

#### SAC wave responses

For measuring SAC dendrite responses to waves, retinas were first bolus loaded with red calcium dye Calbryte^TM^ 590 AM (AAT Bioquest) and then recovered for 30-60 minutes. SACs in the loaded area were then targeted and filled with green calcium dye as described above. Calcium responses from the SAC dendritic field and retinal wave propagation were simultaneously imaged using 950 nm excitation wavelength. Calbryte^TM^ 590 and 520 emissions were separated by photomultiplier tubes filtered for red and green wavelengths, respectively. Images were acquired at 5.92 Hz (128 × 128 pixels, 1ms/line, bidirectional scanning) for 20-30 minutes or at least 30 waves. In some cases, 2 μM gabazine was bath applied to increase the diversity of wave directions. Gabazine application did not affect the strength of tuning of SAC dendrites (Fig S4). To test compartmentalization, SAC responses to waves were recorded for 20-30 minutes in ACSF with no drug, then the same cells were recorded for 20-30 minutes after bath application of 1mM TEA. To measure population SAC activity during cholinergic waves, retinas in P8 ChAT-Cre/nGFP β2-nAChR^−/−^ and littermate controls were bolus loaded with Calbryte 590 AM to measure wave activity. Spontaneous activity was recorded for 20 mins and SAC somas were identified via nGFP expression.

#### SAC morphology

For visualizing P8 SACs morphology, cells in ChAT-Cre/nGFP β2-nAChR^−/−^ and littermate controls were targeted as described above and filled with Alexa 594. Z-stack images were taken using 1040nm excitation at 512×512 pixels, 4 frames per z section. For visualizing P9-11 cell morphology in WT and FRMD7^tm^ mice, average projections of calcium imaging responses were used for tracing. All dendrites were traced using the FIJI/ImageJ Simple Neurite Tracer (SNT) plugin and converted to 2-D skeletons for analysis. Dendritic path length, convex hull area, and Sholl analysis (1μm step length) were computed using traces in SNT. For by quadrant analysis, a MATLAB script segmented skeletons into quadrants which were then auto traced in SNT.

### QUANTIFICATION AND STATISTICAL ANALYSIS

#### SAC quadrant tuning

For light response experiments, raw movies were normalized to ΔF/F0 using FIJI and subsequently imported into MATLAB for further analysis. Light-responsive pixels were identified using a custom MATLAB script that computed the correlation coefficient between each pixel’s ΔF/F0 trace and the light stimulus trace (Fig S1). Pixels exceeding a correlation threshold were classified as responsive and segmented into four quadrants centered on the soma.

For each quadrant, ΔF/F0 responses were averaged across three repetitions of each stimulus direction. Tuning to moving bars was quantified using a direction selectivity index (DSI_light_), defined as the normalized difference between responses to bars moving in the centrifugal (CF) versus centripetal (CP) directions:

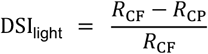

where *R*_CF_ and *R*_CP_ denote the average ΔF/F0 responses to CF and CP bars, respectively.

CF bars were defined as all bars moving outward from the soma within a given quadrant. For example, in the dorsal quadrant (centered at 90°), bars moving toward 45°, 90°, and 135° were classified as CF, while bars moving toward 225°, 270°, and 315° were classified as CP. The cell-level DSI was calculated as the mean DSI across all four quadrants.

For wave imaging experiments, a custom FIJI macro was used to deinterleave red and green channels for separate analysis of retinal waves and SAC dendritic responses, normalize raw movies to ΔF/F0, and identify wave timepoints based on peaks in the average ΔF/F0 wave trace. The propagation direction of each wave was manually scored and binned into eight directions, matching the directions used for the moving-bar stimuli in light response experiments. Wave timepoints, propagation directions, and average quadrant ΔF/F0 responses were then imported into MATLAB for further analysis. To account for variance in SAC activation caused by changes in wave strength, quadrant responses were normalized to the total cell response:

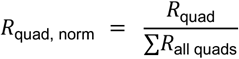

Normalized quadrant responses were used to compute wave DSI:

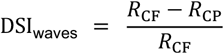

where *R*_CF_ and *R*_CP_ represent the average normalized responses to waves propagating in the CF and CP directions, respectively. CF and CP directions were defined identically to those used for light responses. Cells that did not respond to at least one wave in each quadrant were excluded from analysis.

To quantify tuning as a function of distance from the soma, a custom MATLAB script identified wave-responsive pixels by computing the correlation coefficient between each pixel’s ΔF/F0 trace and the average ΔF/F0 trace of the entire cell. Responsive pixels were binned by quadrant and further subdivided into 10 µm-wide radial bins extending outward from the soma. The radial bins included for analysis were determined by plotting the distribution of wave-responsive pixel counts across bins and excluding distal bins beyond which there was a sharp decline in responsive pixels, indicating little or no dendrites were present. For whole-cell analyses, CF wave preference was computed at each radial step as described above and normalized to the summed responses of the other quadrants at that same step. Only radial steps containing data from all four quadrants were included. For quadrant-specific analyses, when the number of radial bins differed across quadrants, non-normalized ΔF/F0 responses were used to compute DSI_waves_.

#### SAC quadrant wave activation

To measure quadrant-specific wave activation, SAC dendrite calcium responses to retinal waves were recorded in ACSF for 30 minutes. For each cell, wave-evoked responses were summed independently for the nasal, temporal, dorsal, and ventral quadrants. To control for differences in overall response magnitude across cells, quadrant responses were normalized by the total response of the cell. Because normalized quadrant responses constitute compositional data constrained to sum to one, normalized values were transformed using a centered log-ratio (CLR) transform for statistical analysis, which removes the constant-sum constraint and permits valid application of linear models^47^.

#### Statistics

All statistical analyses were performed in MATLAB. For two-sample comparisons, data were first assessed for normality using the Anderson–Darling test. For independent two-sample comparisons, normally distributed data were evaluated using a two-tailed, two-sample Student’s *t*-test, whereas non-normal data were assessed using the non-parametric Mann–Whitney *U* test (Wilcoxon rank-sum test). For paired or within-subject comparisons, normally distributed data were analyzed using a paired *t*-test, and non-normal data were analyzed using the non-parametric Wilcoxon signed-rank test. Summary statistics are reported as mean ± standard deviation unless otherwise noted.

Tuning strength as a function of distance from the soma before and after TEA application was analyzed using a linear mixed-effects model (CF wave preference ∼ drug × distance + (1 | cell)). Drug condition and distance were treated as fixed effects, with cell identity included as a random intercept to account for repeated measurements. Significance of main effects and interactions was assessed using marginal F-tests.

Sholl intersection counts were analyzed using linear mixed-effects models. Genotype and radial distance were included as fixed effects, and litter pair or cell pair and animal were included as random effects. When a significant interaction involving radial distance was detected, follow-up analyses were performed by averaging Sholl intersections within predefined radial bins and analyzing these binned measures using mixed-effects models to localize genotype-dependent effects along the dendritic arbor. This approach avoids pseudoreplication and correctly estimates variability across experimental units^48^.

To assess asymmetries across SAC quadrants and genotypes we fit mixed-design (split-plot) ANOVAs with ‘Quadrant’ as a within-subject repeated-measures factor and ‘Genotype’ as a between-subjects factor. The model included main effects of Quadrant and Genotype as well as their interaction. Significant main effects or interactions were followed by pairwise comparisons corrected using Tukey’s HSD.

To test whether wave-evoked activity differed across quadrants and between genotypes, we performed a mixed-design ANOVA on CLR-transformed normalized quadrant responses with ‘Quadrant’ (dorsal, nasal, temporal, ventral) as a within-subject repeated-measures factor and ‘Genotype’ (wild-type, FRMD7^tm^) as a between-subjects factor. There was a significant main effect of quadrant (F(3,60) = 6.00, p = 0.0012), but no main effect of genotype on overall response magnitude (F(1,20) = 0.18, p = 0.68) and no significant quadrant × genotype interaction (F(3,60) = 1.67, p = 0.18). Despite the absence of an omnibus interaction, post hoc comparisons revealed a robust nasal–temporal asymmetry in WT cells (p = 8.9 × 10−^4^), whereas no significant differences between quadrants were detected in FRMD7^tm^ cells (all p > 0.40), indicating a loss of spatial bias in the mutant. Direct comparisons between genotypes within individual quadrants revealed no significant differences (all p > 0.05), with a trend toward a genotype effect in the temporal quadrant (p = 0.053).

To test whether dendrite length differed in nasal and temporal quadrants and between genotypes, a performed mixed-design ANOVA with Quadrant (nasal, temporal) as a within-cell factor and Genotype (WT, FRMD7^tm^) as a between-cell factor. There was a significant main effect of Quadrant (F(1,27) = 28.81, p = 1.1e-5) and a significant Genotype × Quadrant interaction (F(1,27) = 8.01, p = 0.0087), but no main effect of Genotype (F(1,27) = 0.38, p = 0.54). Tukey HSD post-hoc comparisons showed that WT cells exhibited markedly longer nasal than temporal dendrites (p = 1.5e-6), whereas FRMD7^tm^ cells showed no significant nasal–temporal difference (p = 0.10). Genotype comparisons within quadrants revealed significantly longer temporal dendrites in FRMD7 cells compared to WT (p = 0.022).

---

**Figure S1.**
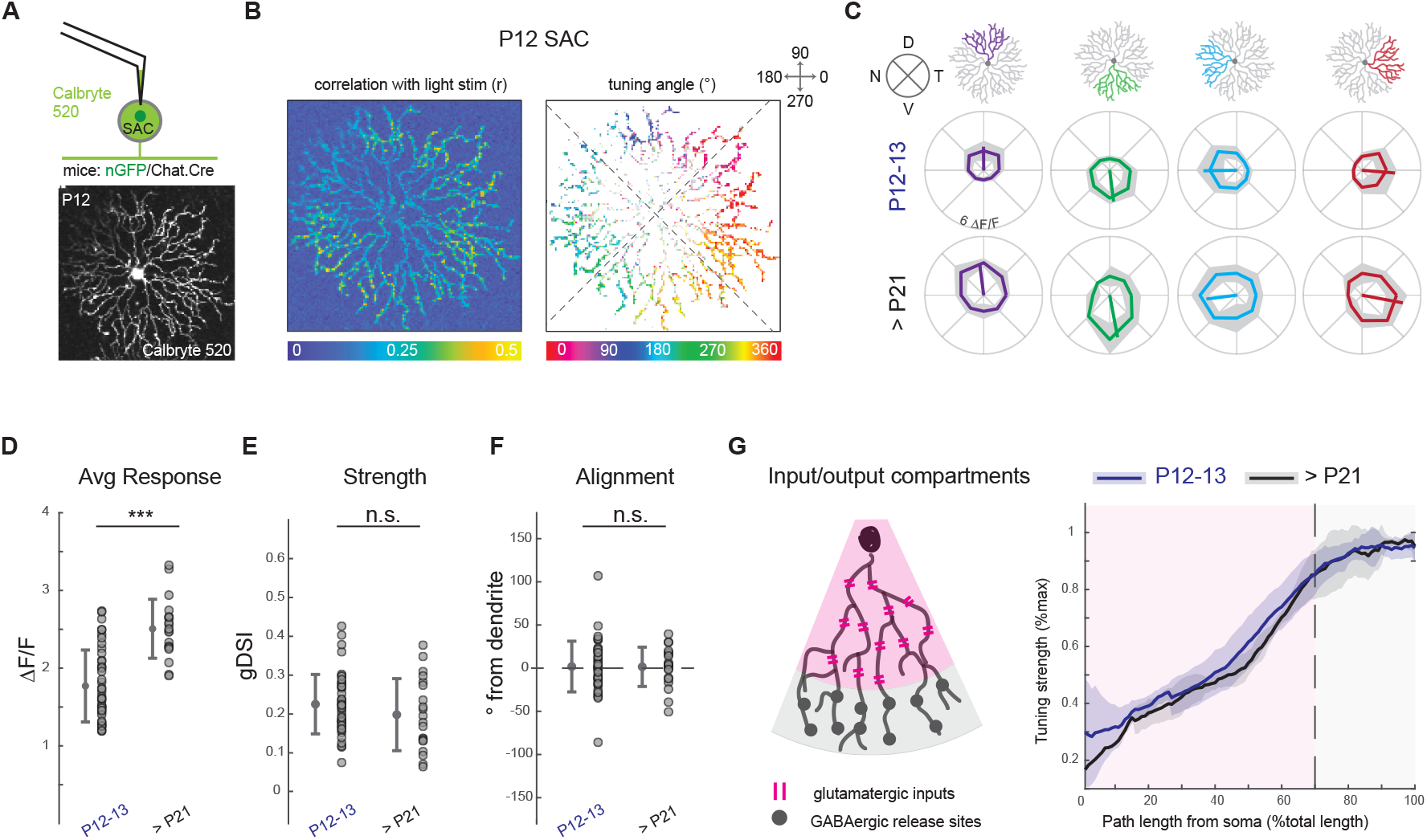
Dendritic direction selective computation is mature prior to eye opening. **(a)** Top: schematic showing targeted sharp bent-tip electrode method of filling SAC dendrites with calcium dye. Bottom: average projection of P12 SAC filled with calcium dye. **(b)** Left: pixel correlation coefficient (r^2^) with light stim trace threshold used to extract light responsive pixels. Right: light-responsive pixels color-coded based on preferred direction (vector sum angle). **(c)** Summary data showing average tuning curves for each quadrant of all cells at P12-P13 (n=10, top) and adult (n = 5, bottom). Lines show vector sum of average response, shaded regions show standard deviation across cells. **(d)** Average magnitude of ΔF/F_0_ for each quadrant regardless of bar direction. Cells in adult show an overall higher response amplitude (two-sample t-test, p = 0.003) (e) No difference across ages in tuning strength for each quadrant as measured by global DSI (gDSI, vector sum normalized by the scalar sum). **(f)** No difference across ages in tuning alignment for each quadrant, measured as the angular difference between the center of the quadrant and the angle of the vector sum of the quadrant tuning curve. **(g)** Left: Schematic showing segmenting of quadrant into input (pink) and output (grey) compartments. Dendrite tuning strength (% max gDSI) as a function of distance from the soma. Individual cell traces (gray) were smoothed with a sliding mean (30 µm window) and normalized to each cell’s maximum; lines represent the adult population mean (black), P12-13 population mean (blue) and ±1 standard deviation (shaded regions). Both ages show similar pattern of increased tuning across the input compartment and a plateau in the output compartment.

**Figure S2.**
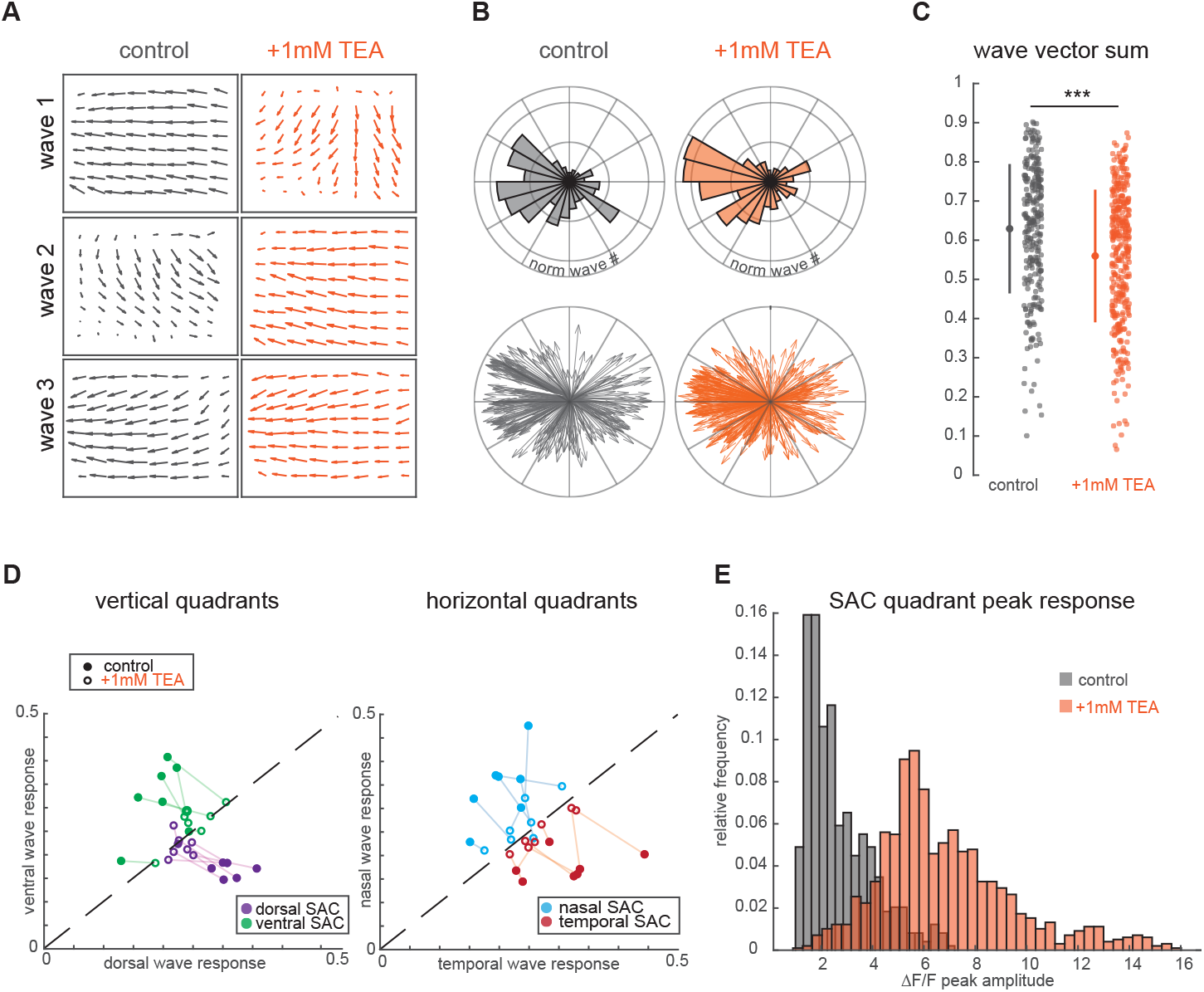
TEA application does not alter directional properties of waves but decreases SAC quadrant direction-selectivity. **(a)** Vector flow fields of three example waves in control conditions (left) and after application of TEA (right). Overall coherence of wave propagation direction is similar across conditions. **(b)** Top: polar histograms showing distributions of wave directions across control (n = 4 FOVs, 262 waves) and TEA conditions (n = 4 FOVs, 345 waves). Bottom: polar plot where each arrow represents a retinal wave, the angle represents that wave’s average direction, and the length represents the strength of the directionality (0–1), where 0 means that the wave had no net direction and 1 means that all pixels in the field of view exhibited the same direction). **(c)** Wave vector sum magnitude across each condition. Waves in TEA show decrease in vector sum (two-sample t-test p = 5.72e-7) though it is a small effect size (Cohen’s d, WT - TEA: 0.415) with waves in TEA still demonstrating strong directionality. **(d)** Left: vertical (dorsal and ventral) quadrants normalized response to ventral waves (y axis) vs dorsal waves (x axis). Each dot is a quadrant, and quadrants are color-coded based on orientation. All ventral quadrants show a preference for ventral waves (dots are above the unity line), while all dorsal quadrants show a preference for dorsal waves (below unity line). Open circles paired with lines represent the same quadrant response after application of TEA. Right: same plot for horizontal quadrants and waves. **(e)** Distribution of peak amplitude of quadrant responses to waves in control (grey) and after application of TEA (orange). TEA application shifts overall response amplitude up, but the long distribution tail indicates sensor is not saturated in the drug condition.

**Figure S3.**
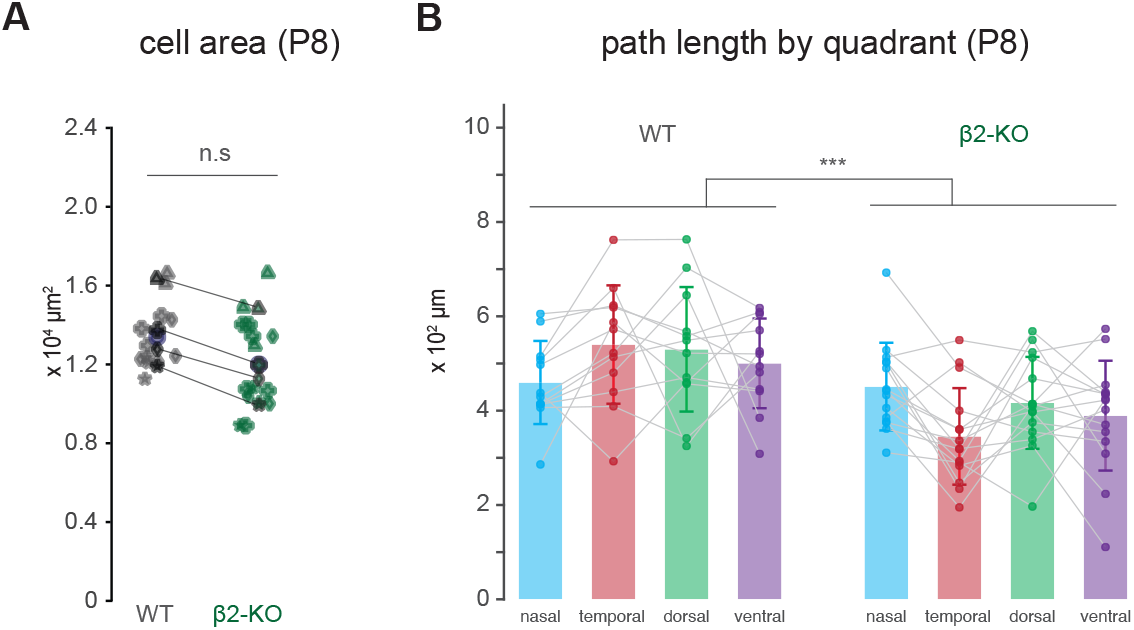
SAC area and path length by quadrant at P8. **(a)** P8 SAC convex hull area of wild-type (n = 12 cells from 4 mice) and β2-nAChR-KO (n = 15 cells from 4 mice) SAC dendrites. Different symbols denote cells from littermate pairs and lines connect littermate averages. No significant difference found in overall cell area, although there was a decreasing trend in littermate averages in β2-nAChR-KO mice. **(b)** Path length by quadrant in wild-type (left) and β2-nAChR-KO (right) mice. A mixed-design ANOVA revealed a significant main effect of genotype (F(1,25)=17.9, p=2.7e-4) and no main effect of quadrant (F(3,75) = 0.54, p = 0.65).

**Figure S4.**
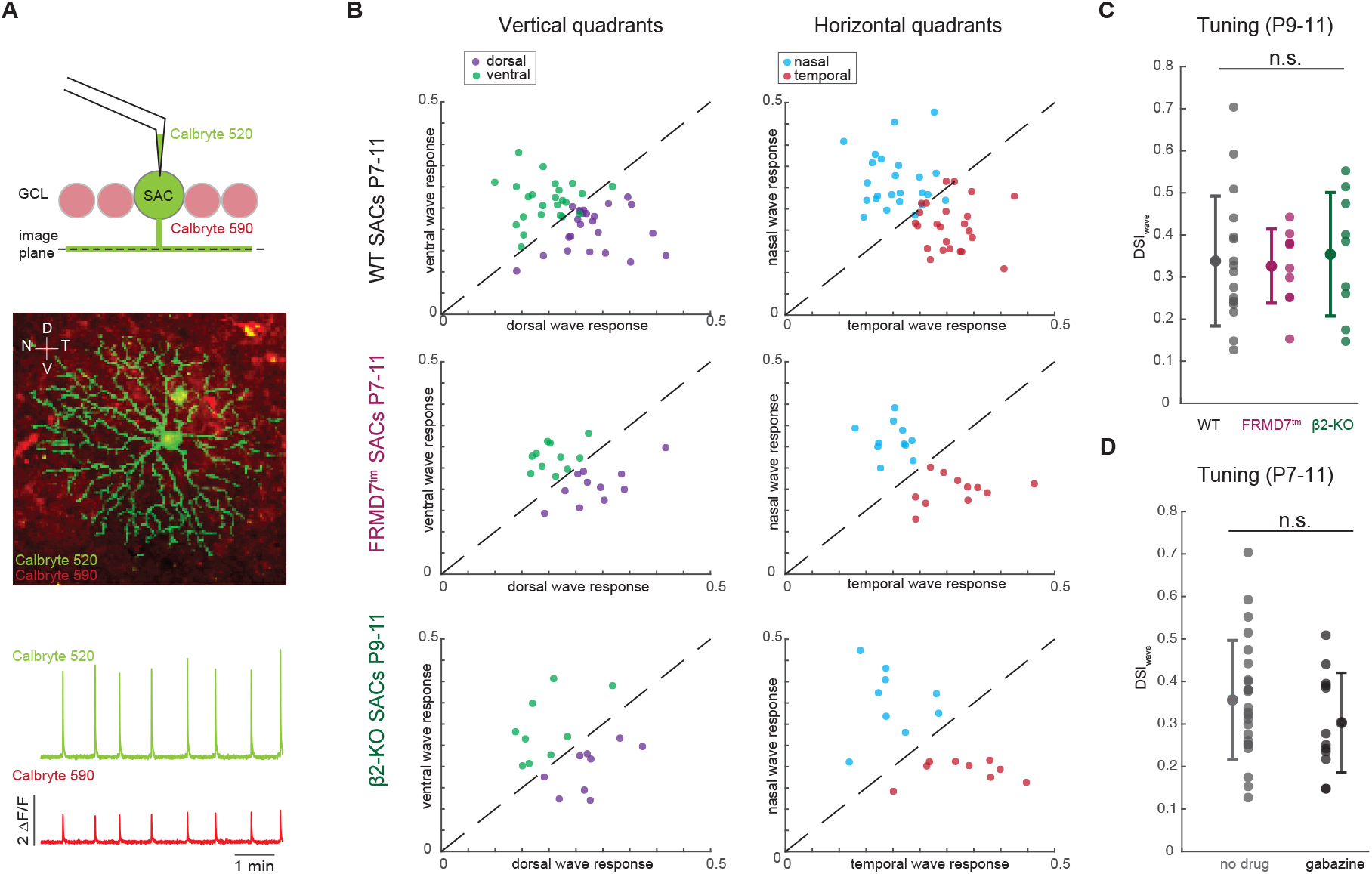
SAC dendrites with altered retinal waves show direction-selective dendritic computation. **(a)** Top: schematic showing dual-colored imaging setup for simultaneous measure of wave propagation (red) and SAC dendrite response (green). Middle: image showing average projections of green and red channel overlayed. Bottom: example whole field ΔF/F traces showing dendrite (green) and wave (red) calcium activity. **(b)** Left: vertical (dorsal and ventral) quadrants normalized response to ventral waves (y axis) vs dorsal waves (x axis). Each dot is a quadrant, and quadrants are color coded based on orientation. All quadrants showing preference for ventral waves are above the unity line while all quadrants showing a preference for dorsal waves are below the unity line. Right: same plot for horizontal quadrants and waves. Rows show the same plots for SACs from WT (n = 24 cells), FRMD7^tm^ (n = 11 cells) and β2-nAChR-KO mice (n = 9 cells). Across all genotypes, quadrants show a preference for waves propagating in their centrifugal direction over the centripetal direction (paired within-cell comparisons; all effects significant after Holm– Bonferroni correction, p≤ 3e-3). **(c)** No difference in SAC tuning strength (DSI_wave_) across genotypes. **(d)** Across all genotypes, no difference found in tuning strength in the absence (n = 30 cells) and presence (n = 14 cells) of 5 µM gabazine.

